# Division of labour within psyllids: Metagenomics reveals an ancient dual endosymbiosis with metabolic complementarity in the genus *Cacopsylla*

**DOI:** 10.1101/2023.04.17.537237

**Authors:** Jessica Dittmer, Erika Corretto, Liliya Štarhová Serbina, Anna Michalik, Eva Nováková, Hannes Schuler

## Abstract

Hemipteran insects are well-known for their ancient associations with beneficial bacterial endosymbionts, particularly nutritional symbionts providing the host with essential nutrients such as amino acids or vitamins lacking from the host’s diet. Thereby, these primary endosymbionts enable the exploitation of nutrient-poor food sources such as plant sap or vertebrate blood. In turn, the strictly host-associated lifestyle strongly impacts the genome evolution of the endosymbionts, resulting in small and degraded genomes. Over time, even the essential nutritional functions can be compromised, leading to the complementation or replacement of an ancient endosymbiont by another, more functionally versatile, bacterium. Herein, we provide evidence for a dual primary endosymbiosis in several psyllid species. Using metagenome sequencing, we produced the complete genome sequences of both the primary endosymbiont ‘*Candidatus* Carsonella ruddii’ and an as yet uncharacterized *Enterobacteriaceae* bacterium from four species of the genus *Cacopsylla*. The latter represents a new psyllid-associated endosymbiont clade for which we propose the name ‘*Candidatus* Psyllophila symbiotica’. Fluorescent *in situ* hybridisation confirmed the co-localization of both endosymbionts in the bacteriome. The metabolic repertoire of *Psyllophila* is highly conserved across host species and complements the tryptophan biosynthesis pathway that is incomplete in the co-occurring *Carsonella.* Unlike co-primary endosymbionts in other insects, the genome of *Psyllophila* is almost as small as the one of *Carsonella*, indicating an ancient co-obligate endosymbiosis rather than a recent association to rescue a degrading primary endosymbiont.

**IMPORTANCE:** Heritable beneficial bacterial endosymbionts have been crucial for the evolutionary success of numerous insects, enabling the exploitation of nutritionally limited food sources such as vertebrate blood and plant sap. Herein, we describe a previously unknown dual endosymbiosis in the psyllid genus *Cacospylla*, consisting in the primary endosymbiont ‘*Candidatus* Carsonella ruddii’ and a co-occurring *Enterobacteriaceae* bacterium for which we propose the name ‘*Candidatus* Psyllophila symbiotica’. Its localization within the bacteriome and its small genome size confirm that *Psyllophila* is a co-primary endosymbiont widespread within the genus *Cacopsylla.* Despite its highly eroded genome, *Psyllophila* complements the tryptophan biosynthesis pathway that is incomplete in the co-occurring *Carsonella.* Moreover, the genome of *Psyllophila* is almost as small as the one of *Carsonella*, indicating an ancient dual endosymbiosis rather than a recent acquisition of a new symbiont. Hence, our results shed light on the dynamic interactions of psyllids and their endosymbionts over evolutionary time.

## INTRODUCTION

Numerous insects maintain long-lasting associations with heritable bacterial endosymbionts that provide the host with essential nutrients lacking from its diet (1). Plant sap-feeding and blood-feeding insects in particular are well-known to harbour nutrient-providing endosymbionts in specialized cells called bacteriocytes, which may form a tissular structure called a bacteriome (2, 3). These so-called primary endosymbionts are obligatory for host survival and reproduction, as they provide essential amino acids and/or vitamins that the host cannot produce or obtain from its food source (4–9). Hence, these bacteria have been crucial for the evolutionary success of numerous insects, enabling the exploitation of nutritionally unbalanced food sources such as vertebrate blood and plant sap.

In turn, the host-associated lifestyle has a strong impact on the genome evolution of the endosymbionts: Their strictly intracellular environment, small effective population size and frequent bottlenecks due to vertical transmission result in genomic decay through the accumulation of deleterious mutations (Muller’s ratchet) and the loss of genes that are no longer needed (10–12). Over evolutionary time, this has produced some of the smallest bacterial genomes known to date (6), streamlined for the production of nutrients required by the host. However, eventually even these pathways can be degraded, leading either to the complementation or the replacement of the ancient endosymbiont by another, more functionally versatile, bacterium (13–18).

This dynamic can be observed in several plant sap-feeding hemipterans which rely on more than one primary endosymbiont to produce all necessary nutrients. Notably, the Auchenorrhyncha (cicadas, planthoppers, spittlebugs) are well-known for their ancient dual endosymbiotic consortia where two co-primary endosymbionts jointly produce the complete set of essential nutrients required by the host, resulting in an intricate metabolic interdependence between the different partners (19). Nonetheless, multiple endosymbiont replacements occurred over time to compensate for the extreme genome erosion of the ancient symbionts (6, 16, 20–25). A similar pattern occurs in aphids (Sternorrhyncha): Whereas most species harbour a single primary endosymbiont, *Buchnera aphidicola*, which provides the host with the ten essential amino acids and the vitamins biotin and riboflavin (26), dual-endosymbiotic systems have evolved repeatedly in multiple aphid lineages to compensate for lost pathways in *B. aphidicola* (13, 14, 18, 27–29).

Similar dual primary endosymbioses may be widespread in psyllids (Hemiptera: Psylloidea), a species-rich group of phloem-feeding jumping plant lice. Like other plant sap-feeding insects, psyllids harbour a bacteriocyte-associated primary endosymbiont (‘*Candidatus* Carsonella ruddii’, hereafter *Carsonella*), which provides the host with essential amino acids (30–33). *Carsonella* is present in all investigated psyllid species and exhibits strict host-symbiont co-divergence, suggesting a single infection of a common ancestor of all extant psyllids (30, 34, 35). Its genome is extremely streamlined and figures among the smallest bacterial genomes known to date (157-175 Kbp) (31). Due to this extreme genome reduction, some *Carsonella* strains are no longer able to produce the full complement of essential amino acids, questioning their ability to fulfil their symbiotic function without compensation from host genes or co-occurring symbiotic bacteria (33, 36).

Additional endosymbionts have indeed been observed to co-inhabit the bacteriome with *Carsonella* in several species (3, 37–39). In these cases, *Carsonella* is located in bacteriocytes surrounding the bacteriome, while a second bacterium occurs in the syncytium at the center of the bacteriome. Importantly, the taxonomy of the co-primary endosymbiont varies depending on the psyllid species: Whereas the syncytium-symbiont (‘Y-symbiont’) of the mulberry psyllid *Anomoneura mori* is an uncharacterized *Enterobacteriaceae* bacterium (*Gammaproteobacteria*) (37) whose symbiotic role is unknown, the citrus psyllid *Diaphorina citri* harbours ‘*Candidatus* Profftella armatura’ (*Betaproteobacteria*). The latter is a defensive and nutritional endosymbiont which produces vitamins, carotenoids and a polyketide toxin, i.e. metabolites that are not provided by *Carsonella* (39, 40). In contrast, the psyllid species *Ctenarytaina eucalypti* and *Heteropsylla cubana* harbour symbionts closely related to the insect endosymbionts ‘*Ca.* Moranella endobia’ and *Sodalis*, whose genomes precisely complement several amino acid biosynthesis pathways missing from the co-occurring *Carsonella* strains (33). Despite typical hallmarks of vertically transmitted intracellular bacteria, the genomes of both endosymbionts are less reduced (>1 Mbp), suggesting a more recent acquisition relative to *Carsonella,* presumably to compensate for lost functions in the latter. In addition, numerous psyllid microbiome studies revealed highly abundant but yet uncharacterized *Enterobacteriaceae* bacteria in diverse species from several psyllid families (41–46), suggesting that dual primary endosymbioses may be more widespread in psyllids than previously thought.

Herein, we aim to elucidate the evolutionary and metabolic relationships between psyllids of the genus *Cacopsylla* (Psyllidae) and their *Enterobacteriaceae* endosymbionts. Many *Cacopsylla* species have indeed been shown to harbour highly abundant *Enterobacteriaceae* endosymbionts that are closely-related to the Y-symbiont co-inhabiting the syncytium of the bacteriome in *A. mori* (42, 45, 46). Furthermore, these symbionts were present in all tested individuals of a given species, suggesting that they may represent co-primary endosymbionts widespread in this genus. In this study, we produced the complete genome sequences of both *Carsonella* and the *Enterobacteriaceae* endosymbionts of four *Cacopsylla* species (*C. melanoneura, C. picta, C. pyri* and *C. pyricola*) known to harbour closely-related endosymbionts from our previous metabarcoding studies (45, 46). Fluorescent *in situ* hybridisation confirmed the co-localization of both endosymbionts in the bacteriome. Comparative genomic analyses revealed that the *Enterobacteriaceae* endosymbionts represent a psyllid-associated clade among other insect endosymbionts. Its genome is almost as small as that of *Carsonella* and complements the tryptophan biosynthesis pathway that is compromised in the co-occurring *Carsonella*.

## RESULTS

### All four *Cacopsylla* species harbour two endosymbionts with tiny genomes

To investigate endosymbiont genetic diversity across different *Cacopsylla* species and genotypes, 12 insect metagenomes were sequenced. These metagenomes encompassed four different host species: *C. melanoneura* and *C. picta,* which complete their development on apple (or hawthorn in the case of *C. melanoneura*) and the pear psyllids *C. pyri* and *C. pyricola.* Multiple metagenomes were sequenced for *C. melanoneura* (N=8) and *C. picta* (N=2), covering different Cytochrome Oxidase I (COI) haplotypes and, in the case of *C. melanoneura,* different regions of origin (Aosta Valley vs. South Tyrol, Italy) and different host plants (apple vs. hawthorn) (Table 1). As expected, the majority of the Nanopore reads belonged to the insect genome and only about 5% of the reads (range: 2.94-9.80%) corresponded to non-host reads. Nonetheless, circular genomes of the primary endosymbiont *Carsonella* could be assembled from all metagenomes (Table 1) with coverages of 85-408x. Genome size ranged from 169,917 to 171,920 bp with 14.98-15.53% GC content, similar to previously sequenced *Carsonella* genomes from other psyllid genera (31, 33, 39, 40). The genomes encoded 182-190 protein-coding genes, one ribosomal rRNA operon and 26-27 tRNAs (Table 1). Synteny and gene content were highly conserved across all genomes, with 161 out of 184 orthogroups (87.5%) shared across all twelve genomes (Fig. 1a, b).

**Fig. 1.**
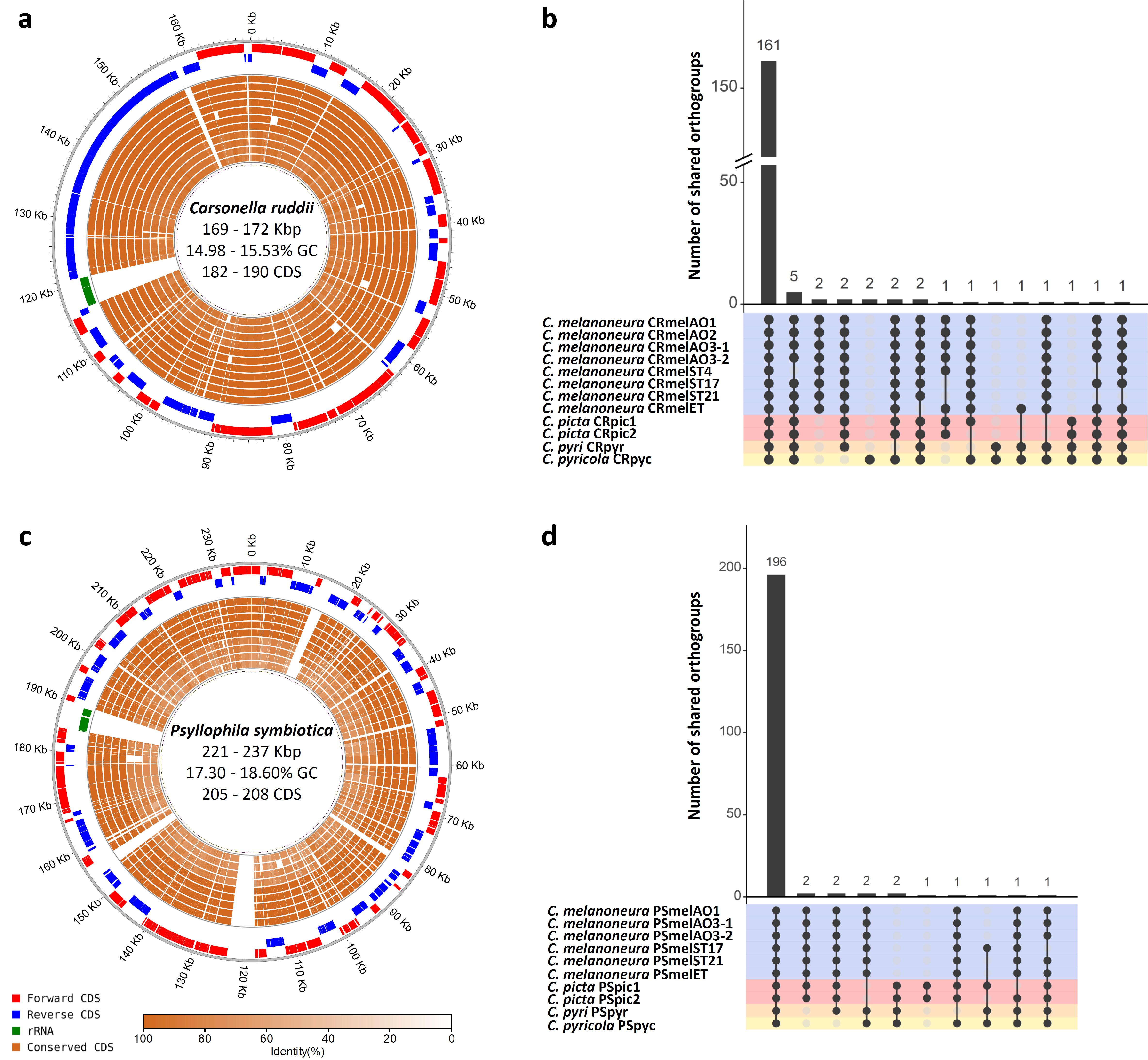
Endosymbiont genomes are highly conserved across *Cacopsylla* host species. (a, c) Circular genome plots of the twelve *Carsonella* genomes (a) and the ten *Psyllophila* genomes (c) produced in this study. The three outer-most circles represent forward CDS, reverse CDS and the ribosomal RNA operon of a reference genome (CRmelAO1 and PSmelAO1, respectively). The inner circles represent the conserved genes in all other genomes of the same taxon (the order is identical to b and d), the shading indicating the degree of sequence similarity compared to the reference genome. (b, d) Intersection plots showing the number of shared orthogroups across all *Carsonella* (b) and *Psyllophila* (d) genomes. The matrix lines are coloured according to host species.

**Table 1.** Properties of the complete endosymbiont genomes obtained in this study. IT=Italy, CZ= Czech Republic. ND=Not determined. Empty columns for Psyllophila indicate that the genome could not be assembled due to insufficient coverage.

| Host species | *Cacopsylla melanoneura* |  |  |  |  |  |  |  | *Cacopsylla picta* |  | *Cacopsylla pyricola* | *Cacopsylla pyri* |
| --- | --- | --- | --- | --- | --- | --- | --- | --- | --- | --- | --- | --- |
| COI haplotype | mel01<sup>a</sup> | mel01<sup>a</sup> | mel01<sup>a</sup> | mel01<sup>a</sup> | mel01<sup>a</sup> | mel01<sup>a</sup> | mel21<sup>b</sup> | mel09<sup>c</sup> | pic18<sup>d</sup> | pic01<sup>e</sup> | pyc01<sup>f</sup> | ND |
| Origin | Aosta Valley (IT) | Aosta Valley (IT) | Aosta Valley (IT) | Aosta Valley (IT) | South Tyrol (IT) | South Tyrol (IT) | South Tyrol (IT) | South Tyrol (IT) | Trentino (IT) | Trentino (IT) | Starý Lískovec (CZ) | Litenčice (CZ) |
| Host plant | Apple | Apple | Hawthorn | Hawthorn | Apple | Apple | Apple | Apple | Apple | Apple | Pear | Pear |
| Sequencing method | Nanopore + Illumina | Nanopore + Illumina | Nanopore + Illumina | Nanopore + Illumina | Nanopore + Illumina | Nanopore + Illumina | Nanopore + Illumina | Nanopore + Illumina | Nanopore + Illumina | Nanopore + Illumina | Nanopore + Illumina | Illumina only |
| *Carsonella* Strain | CRmelAO1 | CRmelAO2 | CRmelAO3-1 | CRmelAO3-2 | CRmelET | CRmelST4 | CRmelST17 | CRmelST21 | CRpic1 | CRpic2 | CRpyc | CRpyr |
| Length (bp) | 169,165 | 169,273 | 169,079 | 169,081 | 169,051 | 169,046 | 169,064 | 169,056 | 171,569 | 171,920 | 168,917 | 169,399 |
| %GC | 15.52 | 15.53 | 15.52 | 15.52 | 15.52 | 15.52 | 15.52 | 15.52 | 15.42 | 15.40 | 14.98 | 15.19 |
| Genes | 216 | 216 | 215 | 215 | 215 | 216 | 215 | 214 | 212 | 211 | 220 | 214 |
| CDS | 186 | 186 | 185 | 185 | 185 | 183 | 185 | 184 | 183 | 182 | 190 | 184 |
| Pseudogenes | 0 | 0 | 0 | 0 | 0 | 3 | 0 | 0 | 0 | 0 | 0 | 0 |
| rRNA | 3 | 3 | 3 | 3 | 3 | 3 | 3 | 3 | 3 | 3 | 3 | 3 |
| tRNA | 27 | 27 | 27 | 27 | 27 | 27 | 27 | 27 | 26 | 26 | 27 | 27 |
| *Psyllophila* strain | PSmelAO1 |  | PSmelAO3-1 | PSmelAO3-2 | PSmelET |  | PSmelST17 | PSmelST21 | PSpic1 | PSpic2 | PSpyc | PSpyr |
| Length (bp) | 237,114 |  | 236,266 | 235,918 | 231,767 |  | 236,072 | 235,744 | 225,730 | 225,865 | 222,755 | 221,413 |
| %GC | 18.26 |  | 18.29 | 18.33 | 18.60 |  | 18.32 | 18.34 | 17.80 | 17.79 | 17.36 | 17.30 |
| Genes | 239 |  | 240 | 239 | 239 |  | 242 | 243 | 240 | 240 | 241 | 240 |
| CDS | 205 |  | 206 | 205 | 205 |  | 207 | 208 | 207 | 207 | 207 | 205 |
| Pseudogenes | 2 |  | 2 | 2 | 2 |  | 3 | 3 | 1 | 1 | 2 | 3 |
| rRNA | 3 |  | 3 | 3 | 3 |  | 3 | 3 | 3 | 3 | 3 | 3 |
| tRNA | 27 |  | 27 | 27 | 27 |  | 27 | 27 | 27 | 27 | 27 | 27 |
| ncRNA | 2 |  | 2 | 2 | 2 |  | 2 | 2 | 2 | 2 | 2 | 2 |

<sup>a</sup> Identical to Accession no. KM206163
<sup>b</sup> Accession no. OQ631077
<sup>c</sup> Identical to Accession no. KM206167
<sup>d</sup> Accession no. OQ631078
<sup>e</sup> Identical to Accession no. KM206174
<sup>f</sup> Accession no. OQ631079

In addition to *Carsonella*, a second circular genome could be assembled from 10 out of the 12 metagenomes (Table 1) with coverages of 19-431x. These genomes belonged to the uncharacterized *Enterobacteriaceae* endosymbiont previously identified through 16S rRNA gene metabarcoding (45, 46). Contigs of this symbiont were also present in the two remaining metagenomes (both from *C. melanoneura*), but the coverage was insufficient to assemble complete genomes. The ten complete chromosomes of the *Enterobacteriaceae* endosymbiont ranged from 221,413 bp in *C. pyri* to 237,114 bp in a strain from *C. melanoneura* from Aosta Valley (strain PSmelAO1, Table 1). GC content varied from 17.30-18.60%. Despite the variations in genome size, synteny and gene content were highly conserved across all *Enterobacteriaceae* genomes (Fig. 1c, d). They contained 205-208 protein-coding genes, 1-3 pseudogenes, one ribosomal rRNA operon, 27 tRNAs and 2 ncRNAs (Table 1). Moreover, 196 out of 209 orthogroups (93.78%) were shared across all ten genomes (Fig. 1d), indicating that the functional repertoire is highly similar across all four host species. Taken together, all four *Cacopsylla* species harbour two endosymbionts with typical hallmarks of a long intracellular symbiotic lifestyle, such as extremely small genomes and low GC content.

### The *Enterobacteriaceae* symbionts represent a new clade of insect endosymbionts

To determine the phylogenetic position of the newly-sequenced psyllid endosymbionts, we performed a Maximum Likelihood phylogenomic analysis based on 67 single-copy genes present in 46 genomes, namely the 10 *Enterobacteriaceae* endosymbionts of *Cacopsylla* spp., 33 insect endosymbionts from the *Gammaproteobacteria* and three *Pseudomonas entomophila* strains as outgroup (Fig. 2). The insect endosymbionts included the two previously sequenced endosymbionts of the psyllid species *C. eucalypti* and *H. cubana* as well as both obligate and facultative endosymbionts of diverse hemipterans (aphids, adelgids, leafhoppers, mealybugs, stinkbugs), beetles (reef beetles and weevils) and the tsetse fly (Table S1). Interestingly, the *Enterobacteriaceae* endosymbionts of *Cacopsylla* spp. were not closely-related to the previously sequenced endosymbionts of the psyllids *C. eucalypti* and *H. cubana* (33) (Fig. 2). Instead, they formed a clade with full bootstrap support that was most closely-related to ‘*Ca.* Annandia adelgestsuga’ and ‘*Ca.* Annandia pinicola’, nutritional endosymbionts of adelgids (47) as well as ‘*Ca.* Nardonella sp.’, ancient endosymbionts of weevils (48) (Fig. 2). Hence, the *Enterobacteriaceae* endosymbionts of *Cacopsylla* spp. represent a new psyllid-associated clade of insect endosymbionts for which we propose the name ‘*Ca.* Psyllophila symbiotica’ (hereafter *Psyllophila*).

**Fig. 2.**
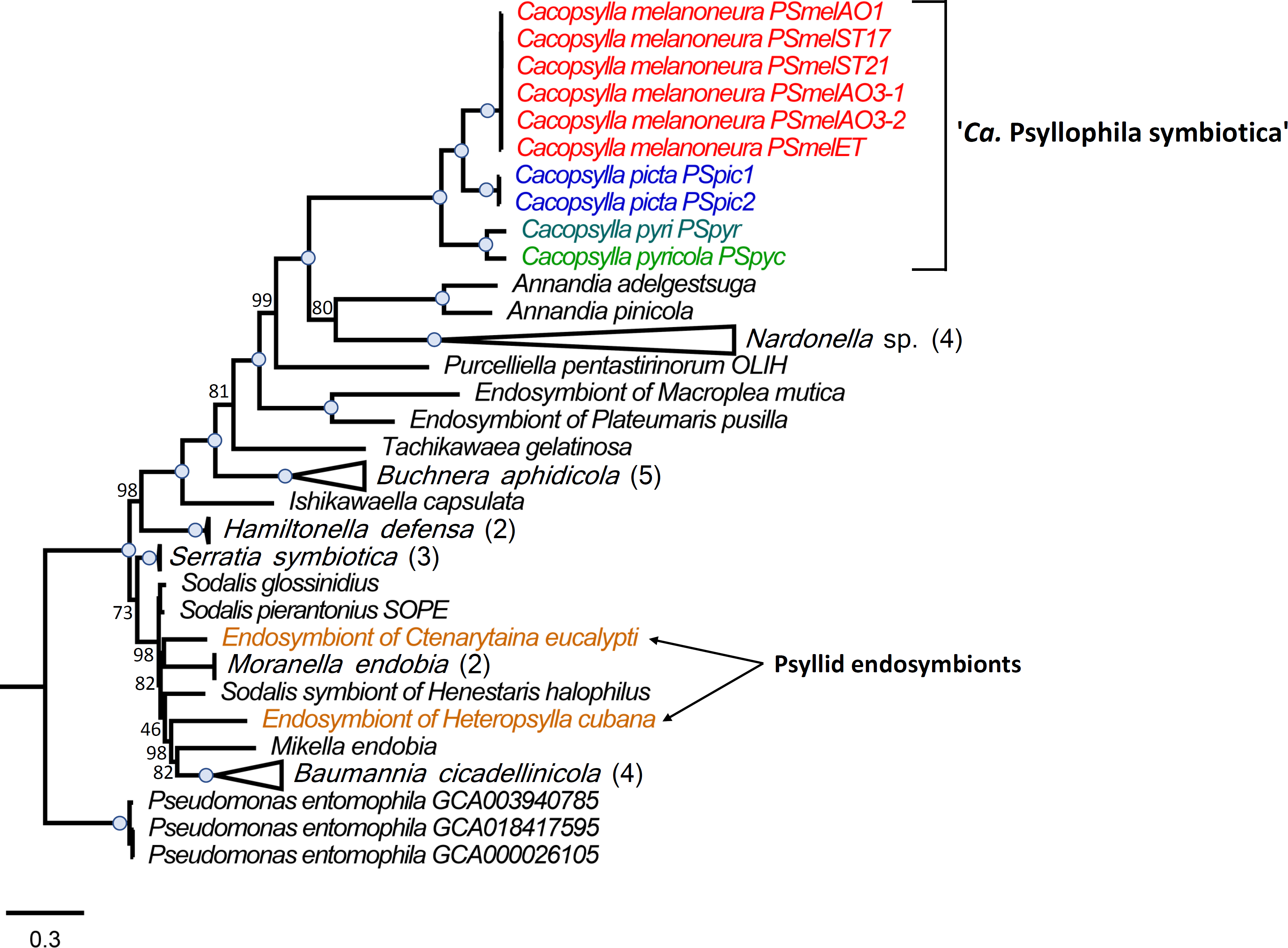
The *Enterobacteriaceae* symbiont represents a new psyllid-associated genus. Maximum-likelihood tree based on the concatenated amino acid sequence alignment of 67 single-copy orthologous genes from 46 genomes, namely the 10 *Enterobacteriaceae* endosymbionts of *Cacopsylla* spp., 33 insect endosymbionts from the *Gammaproteobacteria* and three *Pseudomonas entomophila* strains as outgroup. The *Enterobacteriaceae* endosymbionts of *Cacopsylla* spp. are colour-coded based on host species. Branch support is based on 1000 bootstrap iterations. Blue dots on branches indicate full bootstrap support.

### Both *Cacopsylla* endosymbionts are localized in the bacteriome

Fluorescence *in situ* hybridization with *Carsonella* and *Psyllophila*-specific probes revealed that all *Cacopsylla* species exhibit the same pattern of endosymbiont co-localization in the same bacteriome (Fig. 3). The bacteriomes are large, paired organs localized in the insect’s abdomen. A single bacteriome contains two distinct parts: central and peripheral. Uninucleated bacteriocytes filled with *Carsonella* are located in the peripheral zone of the bacteriome (Fig. 3), whereas the central part is occupied by a multinucleated syncytium filled with *Psyllophila* cells as well as some bacteriocytes containing *Carsonella* (Fig. 3).

**Fig. 3.**
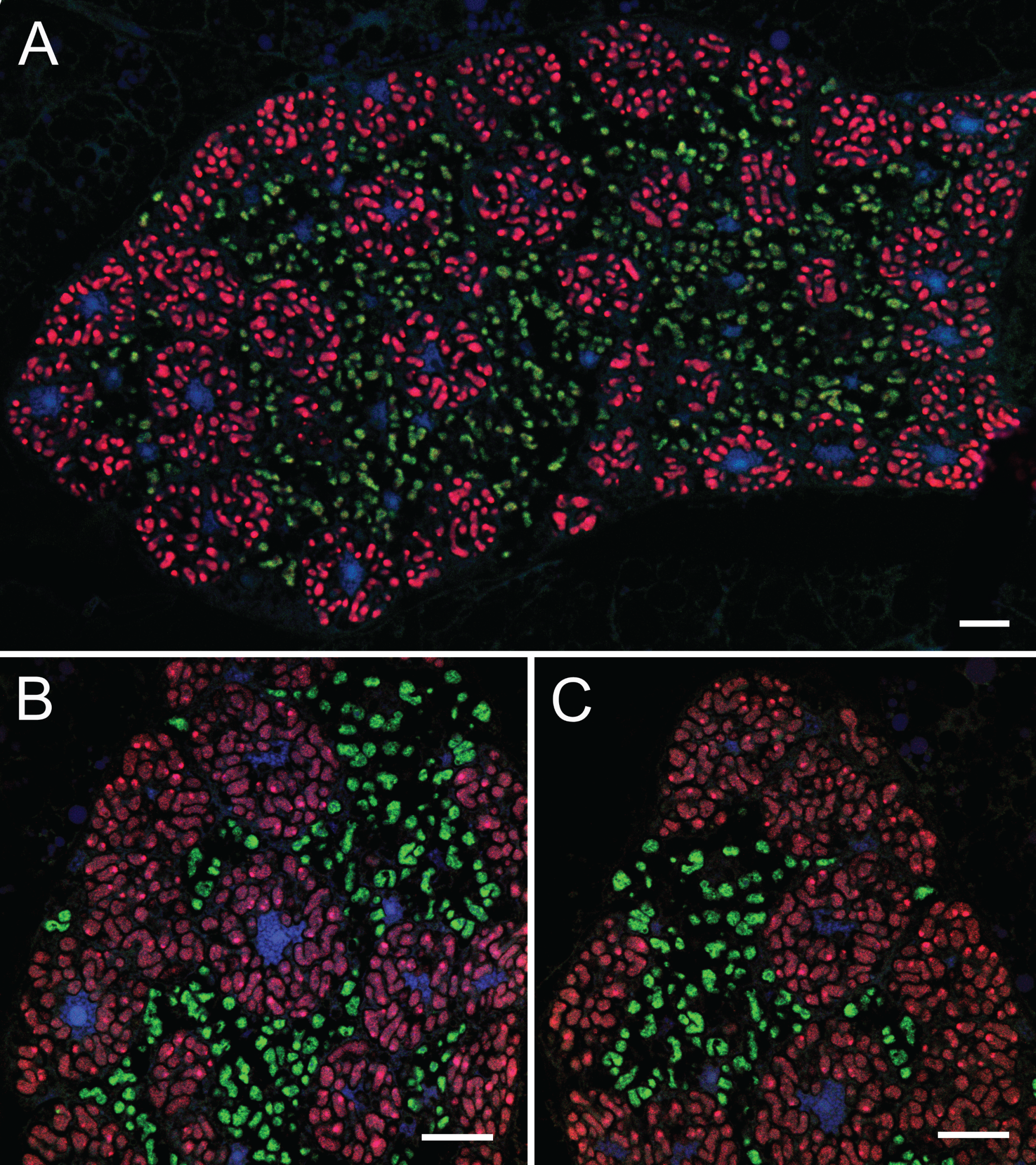
Both *Cacopsylla* endosymbionts are localized in the bacteriome. Fluorescent *in situ* hybridisation of *Carsonella* (Cy3, red) and *Psyllophila* (Cy5, green) symbionts in the bacteriomes of *Cacopsylla pyri* (A, B) and *C. melanoneura* (C). Blue represents DAPI. Scale bar = 10 µm.

### Metabolic complementarity between *Carsonella* and *Psyllophila*

The COG category “Amino acid transport and metabolism” was enriched in all sequenced *Carsonella* genomes (Fig. 4a), in line with its role as a nutritional symbiont. Indeed, based on the KEGG pathway annotation, the biosynthesis pathways for eight of the ten essential amino acids are complete or almost complete in all 12 *Carsonella* strains from the four *Cacopsylla* species (Fig. 4b). Most of the missing functions (*hisN* in the histidine pathway, *dapC* in the lysine pathway, *thrB* in the threonine pathway and *aroE* in the Shikimate pathway, Fig. 4b) are also missing in all previously sequenced *Carsonella* genomes (Table S2, S3). The same applies to the methionine biosynthesis pathway, for which only the last reaction (*metE*) is present in all sequenced *Carsonella* genomes from this and previous studies (Fig. 4b, Table S3). The only difference between the *Cacopsylla*-associated *Carsonella* strains was the absence of *aroB* in the stains from *C. picta* and *C. pyri*, whereas this gene is present in all *Carsonella* strains from *C. melanoneura* and *C. pyricola* (Fig. 4b, Table S3). Interestingly, the tryptophan biosynthesis pathway was incomplete in all 12 *Carsonella* strains from *Cacopsylla* spp., in that only *trpE* and *trpG* were present, whereas the rest of the pathway was missing (Fig. 4b).

**Fig. 4.**
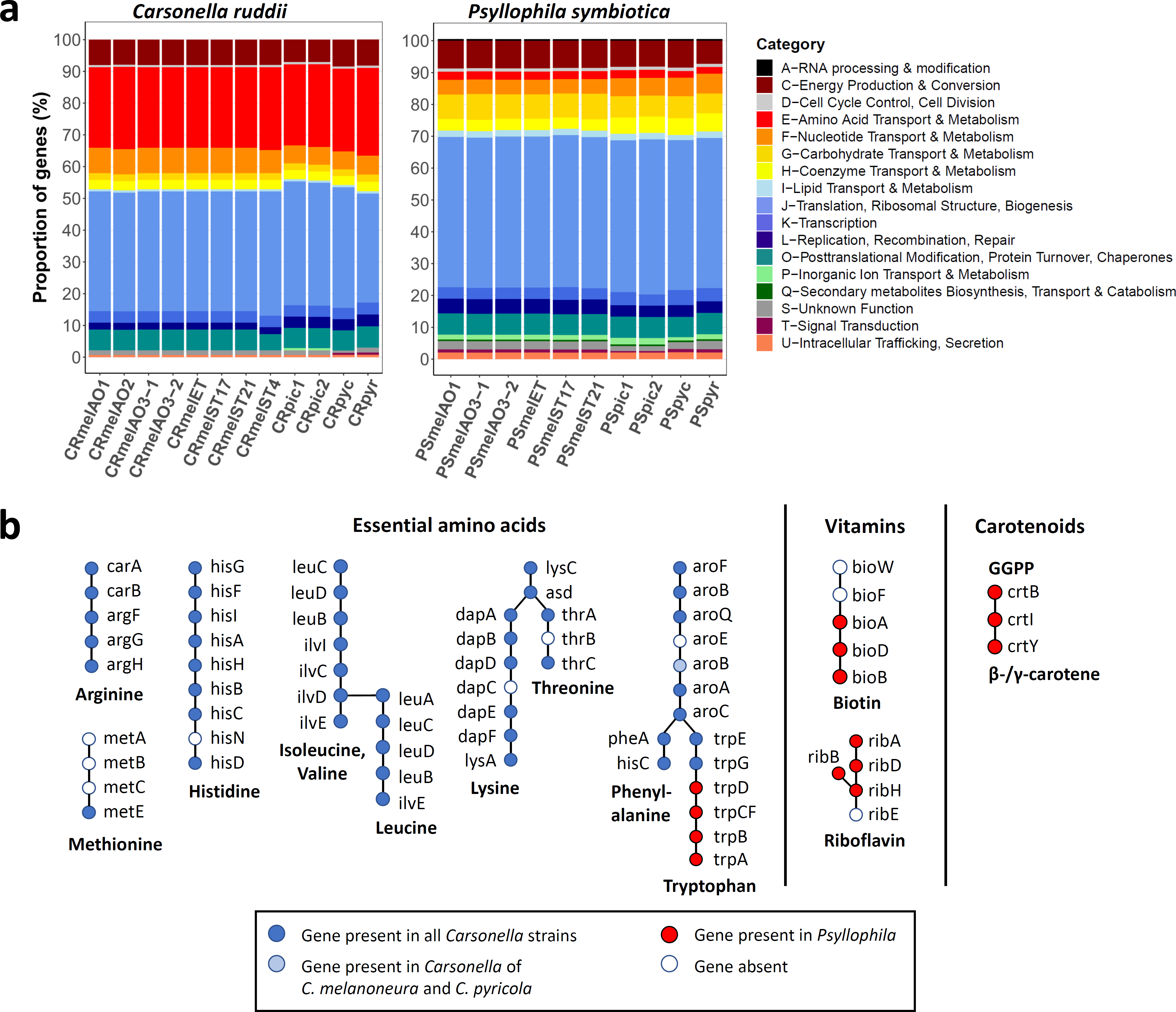
**Metabolic complementarity between *Carsonella* and *Psyllophila.*** (a) COG functional categories for the *Carsonella* and *Psyllophila* genomes show different proportions of genes involved in amino acid transport and metabolism (red). (b) Schematic representation of the metabolic complementarity between the two symbionts for the biosynthesis of essential amino acids, vitamins and carotenoids. Genes present in *Carsonella* genomes are shown in blue, genes present in *Psyllophila* in red.

In contrast to *Carsonella,* the genomes of *Psyllophila* have lost almost all genes involved in amino acid synthesis ((Fig. 4a). Only four genes were retained and these precisely complement the incomplete tryptophan biosynthesis pathway in *Carsonella*, namely *trpD, trpCF, trpB* and *trpA* (Fig. 4b). The four genes were arranged consecutively in the genomes. In addition, all *Psyllophila* genomes encoded partial biosynthesis pathways for the vitamins biotin (*bioA, bioB, bioD*) and riboflavin (*ribA, ribB, ribD, ribH*) as well as all necessary genes for the biosynthesis of carotenoids (*crtB, crtI, crtY*) (Fig. 4b).

### Repeated gene losses throughout *Carsonella* evolution

Apart from the *Carsonella* genomes presented herein, complete genome sequences are available for 11 *Carsonella* strains from nine psyllid species representing five genera and three families (Aphalaridae, Psyllidae and Triozidae) (Table S2). The functional repertoire of these genomes is quite conserved, since 135 out of 197 orthogroups (68.5%) were shared across all 33 genomes and specific orthogroups occurring only in strains from particular host species or genera were rare (14/197) (Fig. 5a). In contrast, host lineage-specific losses of orthogroups were more common. For instance, 12 orthogroups were specifically absent from the three *Carsonella* strains from *H. texana, P. celtidis* and *P. venusta* (Fig. 5a). Similarly, six orthogroups were specifically absent from the *Carsonella* strains from *Ctenarytaina* spp., four orthogroups were absent from strains from *Cacopsylla* spp. and three orthogroups were absent from strains from *Pachypsylla* spp. (Fig. 5a). These differences are also reflected in repeated losses of genes or entirely pathways involved in essential amino acid biosynthesis across the *Carsonella* phylogeny (Fig. 5b). Notably, the tryptophan pathway has been lost at least three times independently, as it is incomplete or missing in all *Carsonella* strains associated with the genera *Cacopsylla* and *Heteropsylla* (Psyllidae) as well as *Ctenarytaina* and *Pachypsylla* (Aphalaridae) (Fig. 5b, Table S3). In contrast, this pathway is complete in the *Carsonella* strains from *Bactericera* spp. (Triozidae) and *Diaphorina citri* (Liviidae) (Fig. 5b, Table S3). Other repeatedly lost functions include the histidine biosynthesis pathway as well as the genes *aroB* and *dapE,* implicated in the Shikimate and lysine pathways, respectively (Fig. 5b, Table S3). Concomitantly, co-primary endosymbionts complementing the missing amino acid biosynthesis pathways have been identified in several species, i.e. complementing tryptophan in *Cacopsylla* spp. (this study) and *H. cubana* (33) and both tryptophan and arginine in *Ct. eucalypti* (33) (Fig. 5b).

**Fig. 5.**
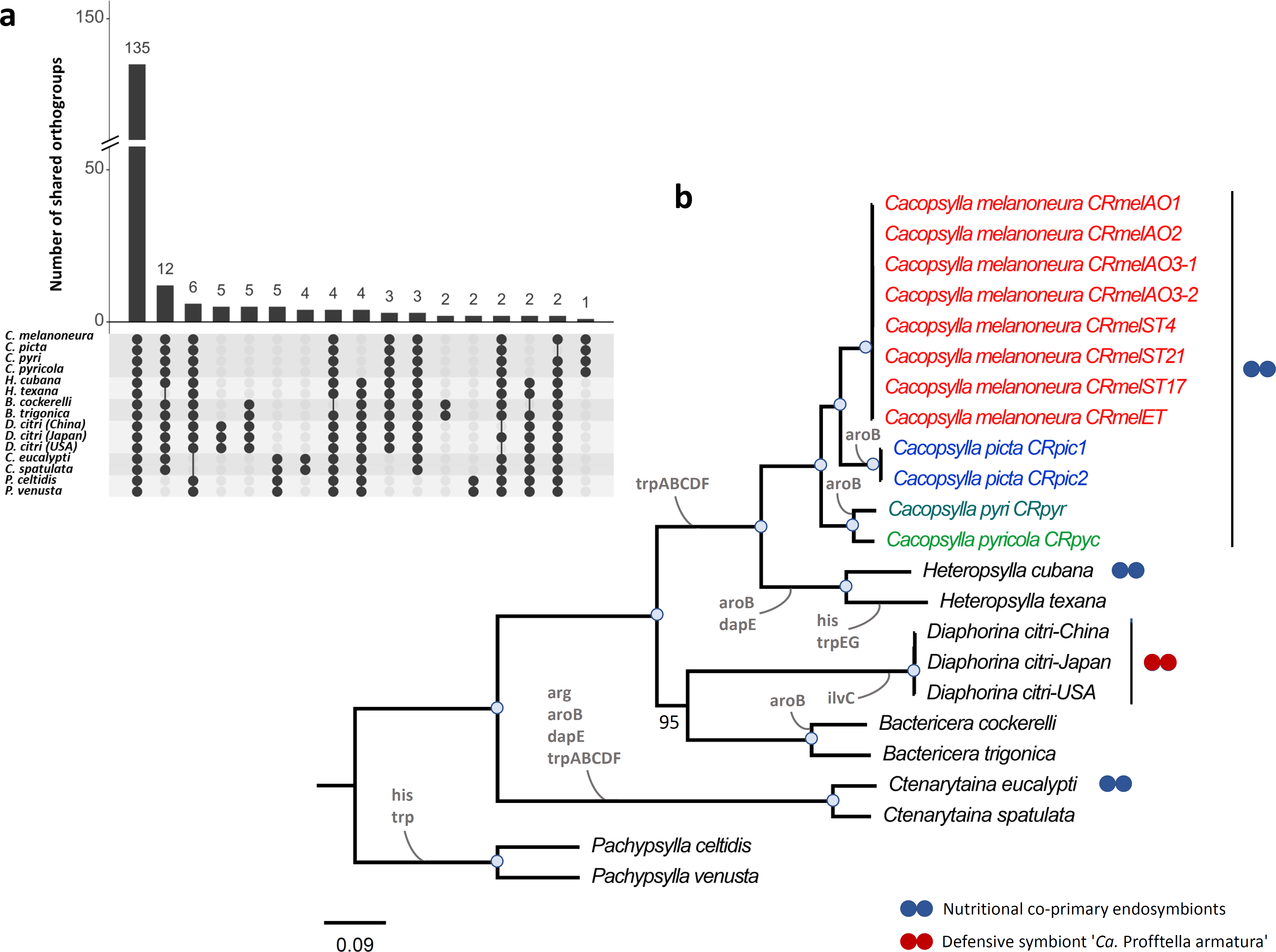
Repeated gene losses throughout *Carsonella* evolution. (a) Intersection plot showing the distribution of orthogroups across 33 *Carsonella* genomes depending on host species. (b) Maximum-likelihood tree based on the concatenated amino acid sequence alignment of 119 single-copy orthologous genes present in all 33 *Carsonella* genomes. The *Carsonella* strains of *Cacopsylla* spp. are colour-coded based on host species. Branch support is based on 1000 bootstrap iterations. Blue dots on branches indicate full bootstrap support. Losses of genes or pathways involved in the biosynthesis of essential amino acids are indicated on the branches. (See Table S3 for a detailed list of genes identified based on KEGG pathway annotation). The presence of known co-primary endosymbionts is indicated using blue dots for amino acid-providing nutritional co-primary endosymbionts and using red dots for the defensive and nutritional endosymbiont ‘*Ca.* Profftella armatura’.

## DISCUSSION

Herein, we present the complete genome sequences of both *Carsonella* and the uncharacterized *Enterobacteriaceae* endosymbionts of four *Cacopsylla* species from different host plants. The *Enterobacteriaceae* endosymbionts represent a psyllid-associated clade among other insect endosymbionts, for which we propose the name ‘*Ca.* Psyllophila symbiotica’. Both endosymbionts co-occur within the bacteriome, exhibiting the same co-localization pattern (*Carsonella* in peripheral bacteriocytes, *Psyllophila* in the central syncytium) as for other dual endosymbioses in *D. citri* and *A. mori* (37–39). In combination with a small and AT-rich genome, the bacteriome localization confirms that *Psyllophila* is a co-primary endosymbiont widespread within the genus *Cacopsylla.* Interestingly, unlike co-occurring endosymbionts in other psyllid species (33, 39, 40), the *Psyllophila* genome is almost as small as the genome of *Carsonella*, indicating an ancient dual endosymbiosis rather than a recent acquisition of a more versatile symbiont to rescue a degrading primary endosymbiont.

Despite having a tiny and functionally limited genome, *Psyllophila* has retained the necessary genes to complement the tryptophan biosynthesis pathway that is compromised in the co-occurring *Carsonella.* This appears to be a recurring theme across *Carsonella* evolution, since the tryptophan pathway is the most frequently lost amino acid biosynthesis pathway based on the genomes available to date. Specifically, this pathway has been lost multiple times independently, namely in the *Carsonella* strains associated with species from the genera *Pachypsylla* and *Ctenarytaina* (Aphalaridae), and the psyllid lineage leading to both *Heteropsylla* and *Cacopsylla* (Psyllidae). Apart from tryptophan, the arginine and histidine pathways have also been lost in specific *Carsonella* strains, albeit less frequently (33). Concomitantly, co-primary endosymbionts complementing the missing amino acid biosynthesis pathways have been identified in several species, namely *Psyllophila* and a *Sodalis*-like symbiont complementing tryptophan in *Cacopsylla* spp. (this study) and *H. cubana* (33), respectively, and another *Sodalis*-like symbiont complementing both tryptophan and arginine in *Ct. eucalypti* (33). Intriguing cases in this context are the psyllid species *H. texana, P. celtidis* and *P. venusta*, whose *Carsonella* strains have lost both the histidine and tryptophan pathways, but no co-primary endosymbionts have been observed to date (33). Possible alternative scenarios are that the missing genes are encoded by the host, e.g. after horizontal transfers of bacterial genes to the host genome, or that the amino acids in question are present in sufficient quantities in the phloem sap of the psyllid’s host plant. Although numerous genes of bacterial origin have indeed been identified in the genome of *P. venusta*, they do not restore the missing amino acid pathways (36).

Based on its genome sequence, *Psyllophila* not only rescues tryptophan biosynthesis, it also encodes partial biosynthesis pathways for the vitamins biotin and riboflavin, as well as all necessary genes for the synthesis of carotenoids, pigments that may protect against oxidative damage of DNA (49). This represents a striking convergence with ‘*Ca.* Profftella armatura’, the co-primary endosymbiont in several *Diaphorina* species, which has an almost identical gene set for these pathways as *Psyllophila* (40). As in *Psyllophila,* the last step in the riboflavin pathway is missing in ‘*Ca.* Profftella armatura’, but the relevant gene has been detected in the genomes of the psyllids *D. citri* and *P. venusta*, likely due to a horizontal transfer from an unknown bacterium (36). Likewise, we identified a similar riboflavin synthase gene encoded in the genomes of all four *Cacopsylla* species via blast searches of the *D. citri* gene against preliminary assemblies of the insect genomes from our metagenomic datasets. Hence, it is likely that riboflavin can be jointly synthesized by *Psyllophila* and its psyllid hosts, just like in the symbiosis of *D. citri* and *Profftella*. In any case, the functional similarity between two distantly-related psyllid endosymbionts highlights the importance of these metabolites for the psyllid hosts and/or the endosymbionts.

Taken together, our data shed light on the dynamic interactions of psyllids and their endosymbionts over evolutionary time. Notably, the tiny and highly eroded genome of *Psyllophila* suggests a long-lasting dual endosymbiosis of *Carsonella* and *Psyllophila* within the genus *Cacopsylla*. However, this dual endosymbiosis has likely reached a highly labile state, since no functional redundancy exists between the two endosymbionts and any additional gene loss would destabilise the symbiotic system. Considering the diversity of predominant psyllid-associated bacteria revealed by previous studies (34, 41–46, 50), it is likely that the ancient endosymbiont *Psyllophila* has already been replaced by younger symbionts in some psyllid lineages. For instance, this may have been the case in *H. cubana* and *C. eucalypti*, which harbour more recently acquired co-primary endosymbionts with larger genomes (33). It is tempting to speculate that species which do not harbour co-primary endosymbionts today (e.g. *P. venusta*) may have harboured a similar dual endosymbiosis in the past, but the co-symbiont was lost without replacement, maybe because its functions were no longer required after a change in ecological conditions (e.g. change of host plant, evolution of gall-forming behaviour). This could also explain why the *Carsonella* strains in these species have lost similar genes and pathways as the strains existing in dual endosymbiotic systems today.

This raises the question whether these pathways were lost before or after the establishment of the dual primary endosymbiosis. According to the Black Queen Theory on the evolution of dependencies within bacterial communities (51), it is advantageous for bacteria to lose costly metabolic functions (i.e. to streamline their genomes), as long as another species within the community still produces these metabolites as “common goods”. Applying this concept to a community with two partners would imply that any essential pathway can be lost in only one of them but has to be retained in the other, to maintain all essential functions in the system. Hence, in the dual endosymbioses in psyllids, *Carsonella* may have lost the tryptophan pathway, since it was encoded by its symbiotic partner. In turn, the co-primary endosymbionts lost all other genes involved in amino acid biosynthesis, since these were maintained in *Carsonella*, thus establishing the existing metabolic complementarities in different psyllids. However, only a few psyllid endosymbionts have been characterized at the genomic level and more studies across the psyllid tree of life will be necessary to obtain a more complete picture of the evolutionary dynamics of psyllids and their primary endosymbionts.

## MATERIALS AND METHODS

### Psyllid samples

Genomic data was obtained from four psyllid species: the apple psyllids *Cacopsylla melanoneura* and *C. picta* as well as the pear psyllids *C. pyri* and *C. pyricola*. All four species are vectors of plant pathogens, namely ‘*Ca.* Phytoplasma mali’ and ‘*Ca.* Phytoplasma pyri’, respectively causing Apple proliferation and Pear decline (52, 53). Remigrants (i.e. adults that return to their host plants for reproduction after overwintering on shelter plants) of *C. melanoneura* were captured in various apple orchards in two Italian regions (Aosta Valley and South Tyrol) in March 2020 and March 2021. Additional *C. melanoneura* specimens were sampled on hawthorn (*Crataegus* sp.) in a single location (Aosta Valley) in March 2021. Remigrants of *C. picta* were captured from apple orchards in Trentino (Italy) in April 2021. Adults of *C. pyri* and *C. pyricola* were collected in pear orchards in Litenčice (Czech Republic) in December 2019 and in Starý Lískovec (Czech Republic) in July 2020, respectively. Sampling was done using the beating tray method.

Since psyllids are too small to obtain sufficient DNA for long-read sequencing from a single individual, several specimens need to be pooled, which introduces genetic variation that can hinder genome assembly. We used two different strategies to reduce the genetic variation among the pooled individuals, depending on the host plant of the different species. For *C. melanoneura* and *C. picta* collected on apple trees, we applied the same experimental design as in (52): In the green house, the field-caught adults were sorted into mating couples and each couple was caged on a branch of an apple tree (cultivar Golden Delicious) using nylon nets. Once the offspring of the mating couples had reached adulthood, all newly-emerged siblings were collected and stored at -20°C. Since the primary endosymbionts are vertically transmitted from mother to offspring, all siblings harbour genetically identical endosymbionts and can therefore be pooled without introducing genetic variation for the endosymbionts. In addition, the Cytochrome Oxidase I (COI) haplotype was determined for two individuals per sibling group according to (54), to determine the genetic diversity among the different populations and mating couples. For all psyllids that do not develop on apple (*C. melanoneura* from hawthorn and the pear psyllids *C. pyri* and *C. pyricola*), adults collected in the field were immediately stored at -20°C. Subsequently, the COI haplotype was determined for numerous individuals of each species, in order to select individuals with identical COI haplotype for pooling.

### DNA extraction

For each sibling group of the apple psyllids *C. melanoneura* and *C. picta* selected for long-read metagenome sequencing, DNA was extracted from two pools, each containing 4-6 whole females. For the field-caught *C. melanoneura* from hawthorn and *C. pyricola* from pears, DNA was first extracted from individual females and subjected to COI haplotype determination as outlined above. Subsequently, two pools, each combining the DNA extracts of five females with identical haplotypes, were established for each species. Only females were used since they are larger and hence provide more DNA and because we reasoned that their endosymbiont titers may be higher since the endosymbionts are harboured in two tissues, the bacteriome and the ovaries. DNA extraction was performed using a modified protocol of the PureGene Tissue kit (Qiagen). Whole insects were ground in 100 µl of Cell Lysis Solution and 5 µl Proteinase K solution and incubated at 56°C for three hours, followed by an incubation with 1.5 µl RNase A at 37°C for 30 minutes. Subsequently, proteins were precipitated by adding 35 µl of Protein Precipitation Solution. DNA was then extracted with one volume chloroform/isoamyl alcohol (24:1 v/v) and precipitated in one volume of isopropanol after overnight incubation at -20°C. The DNA pellet was resuspended in 40 µl of sterile water and incubated at 65°C for one hour to increase DNA rehydration. For *C. pyri,* DNA was extracted from a single female using the QIAamp DNA Micro Kit (Qiagen) according to the manufacturer’s instructions.

### Metagenome sequencing and assembly

Long-read metagenome sequencing using an Oxford Nanopore-Illumina hybrid approach was performed for *C. melanoneura, C. picta* and *C. pyricola*. For each sample, one pool was used for long-read sequencing on the MinION (Oxford Nanopore Technologies, UK) and the second pool was used for 2 × 150 bp paired-end sequencing on an Illumina NovaSeq (Macrogen). About 1.5 µg of DNA was used for library preparation using the Oxford Nanopore Ligation Sequencing kit SQK-LSK 109 (Oxford Nanopore Technologies, UK). Each library was sequenced on an entire R9.4 flowcell for 43-72 hours, depending on pore activity. Basecalling was done using Guppy v5.0.11 (Oxford Nanopore Technologies, UK) in high-accuracy mode. Low quality (< Q7) and short (< 500 bp) reads were discarded and host reads were removed via mapping against a genome scaffold of *C. melanoneura* (J. M. Howie & O. Rota-Stabelli, unpublished data) using Minimap2 v2.15 (55). The remaining non-host reads ≥ 500 bp were assembled using Flye v2.9 (56) with the --metagenome option. Contigs belonging to the endosymbionts were identified using blast (57). Reads were mapped back onto the endosymbiont contigs using Minimap2 v2.15 and all mapped reads were assembled again with Flye v2.9 using the same parameters. This produced two circular genomes for most datasets. These genomes were first polished with Nanopore reads using Medaka v1.5.0 (https://github.com/nanoporetech/medaka) and subsequently with Illumina reads using several iterations of Polca, a genome polisher integrated in the MaSuRCa toolkit v4.0.7 (58), until no more errors were found. It is important to note that the two endosymbiont genomes need to be polished together to avoid the introduction of errors in highly conserved regions (e.g. ribosomal RNA operon) of the endosymbiont genome with lower coverage. In rare cases, two rounds of Flye assemblies did not produce circular endosymbiont genomes. Two of these genomes (CRmelAO2 and PSmelET) could be finished using alternative assembly approaches: (i) Assembly with Canu v2.1.1 (59), polishing with Medaka and Polca as outlined above, followed by scaffolding and gap-closing with Redundans v0.14 (60) and (ii) Nanopore and Illumina reads mapping onto the complete endosymbiont genomes were assembled together using SPAdes v3.15.1 (61). Genome coverage was estimated by mapping the Nanopore reads onto the finished genomes during the polishing step with Medaka.

The metagenome of *C. pyri* was assembled from 42 million 2 × 250 paired-end reads from a single female sequenced on an Illumina NovaSeq (University of Illinois, Urbana-Champaign, USA). The metagenome was assembled using SPAdes v3.15.1 (61) with the --meta option and default kmers. Endosymbiont contigs were identified based on coverage, which initially produced two contigs for *Carsonella* and three contigs for the *Enterobacteriaceae* symbiont. These contigs were ordered based on the complete genomes obtained using long-read sequencing and closed after scaffolding and gap-closing with Redundans v0.14 (60). The completeness of all genomes was assessed using BUSCO (gammaproteobacteria_odb10 dataset) (62).

### Functional genome analyses

All complete endosymbiont genomes were annotated using the NCBI Prokaryotic Genome Annotation Pipeline (PGAP) version 2021-07-01.build5508 (63). Circular plots of conserved protein-coding genes were produced using MGCplotter (https://github.com/moshi4/MGCplotter). For each endosymbiont (i.e. *Carsonella ruddii* and *Psyllophila symbiotica*), all protein-coding genes were assigned to orthogroups using Orthofinder v2.5.2 (64) and shared orthogroups were plotted using the package UpsetR (65). Clusters of Orthologous Genes (COG) categories were determined using eggNOG-mapper v2.1.7 (66) and KEGG pathway annotations were obtained using BlastKOALA v2.2 (67).

### Phylogenomics

Orthofinder v2.5.2 (64) was used to identify single-copy orthologous genes shared (i) between all available *Carsonella* genomes and (ii) between the *Enterobacteriaceae* endosymbionts of *Cacopsylla* spp., 33 other nutritional endosymbionts from the *Gammaproteobacteria* and three *Pseudomonas entomophila* strains as outgroup. The amino acid sequences of each conserved gene were aligned using Muscle v3.1.31 (68) and the alignments were concatenated into a partitioned supermatrix using the script geneStitcher.py (https://github.com/ballesterus/Utensils/blob/master/geneStitcher.py). IQ-TREE v1.6.1 (69) was used to predict the optimal amino acid substitution model for each gene partition (70, 71) and to produce a Maximum Likelihood phylogenetic tree with 1000 bootstrap iterations. The tree was visualized in FigTree v1.4.4 (https://github.com/rambaut/figtree).

### Fluorescence *in situ* hybridisation

Fluorescence *in situ* hybridization was conducted with symbiont-specific probes complementary to their 16S rRNA gene sequences (*Carsonella*: Probe Carso107 ‘Cy3-ATACTAAAAGGCAGATTCTTG’, *Psyllophila*: Probe Psyllo118 ‘Cy5-TCCATTGAGTAGTTTCCCAG’). For *Carsonella*, two helper probes were used to increase the signal (helperFCarso107: ‘AGCGAACGGGTGAGTAATATG’, helperRCarso107: ‘ACATTTCTATATACTTTCCA’).

Insects preserved in ethanol were rehydrated and then postfixed in 4% paraformaldehyde for two hours at room temperature. Next, the specimens were dehydrated again by incubation in increased concentrations of ethanol and acetone, embedded in Technovit 8100 resin (Kulzer, Wehrheim, Germany), and cut into semithin sections (1 µm). The sections were then incubated overnight at room temperature in hybridization buffer containing the specific probes at a final concentration of 100 nM. After hybridization, the slides were washed three times in PBS, dried, covered with ProLong Gold Antifade Reagent (Life Technologies) and observed using a Zeiss LSM 900 Airyscan 2 confocal laser scanning microscope.

## DATA AVAILABILITY

The genomes produced in this work are accessible in the NCBI database under BioProject accessions PRJNA803426 (endosymbionts of *Cacopsylla melanoneura*), PRJNA853274 (endosymbionts of *Cacopsylla picta*), PRJNA853282 (endosymbionts of *Cacopsylla pyricola*) and PRJNA853726 (endosymbionts of *Cacopsylla pyri*). New COI haplotypes for *C. melanoneura, C. picta* and *C. pyricola* were deposited under accessions OQ106377 – OQ106379.

## ACKNOWLEDGEMENTS

We thank Domagoj Gajski, Michal Štarha and Valentina Candian for help collecting psyllids in the field as well as Massimiliano Trenti for help with the psyllid rearing in the greenhouse. This study was funded by a joint project of the province of Bolzano and the Austrian Science Fund FWF (FIGHToplasma, I 4639-B). Anna Michalik was supported by the Polish National Science Centre grant 2017/26/D/NZ8/00799.

## SUPPLEMENTARY MATERIAL

**Supplementary Table S1.** List of insect endosymbiont genomes included in the phylogenomics analysis of ‘*Ca*. Psyllophila symbiotica’.

**Supplementary Table S2.** Table of previously published *Carsonella* genomes used for comparative and phylogenomic analyses.

**Supplementary Table S3.** Presence of genes involved in the biosynthesis of essential amino acids in all available *Carsonella* genomes based on KEGG pathway annotations using BlastKoala.

